# MSIanalyzer: Targeted Nanopore Sequencing Enables Single Nucleotide Resolution Analysis of Microsatellite Instability Diversity

**DOI:** 10.1101/2025.06.26.661510

**Authors:** Ting Zhai, Daniel J. Laverty, Zachary D. Nagel

## Abstract

We present a targeted sequencing-based pipeline that profiles microsatellite instability (MSI) at single-nucleotide resolution. Targeted amplicons from the five widely studied Bethesda panel microsatellite loci were sequenced using Oxford Nanopore Technology in two microsatellite unstable colorectal cancer cell lines (HCT15, HCT116), two microsatellite stable cancer cell lines (TK6, U2OS), and two peripheral blood mononuclear cell samples from healthy donors. An anchor-extension algorithm was developed to capture repeat motifs while allowing interruptions, using a threshold informed by platform-specific error. Cluster-aware Dirichlet-multinomial and beta-binomial tests were applied for between-sample comparisons while accounting for read-level clustering within samples. The algorithm revealed distinct repeat profiles in HCT15 and HCT116 compared to other cell types and uncovered allelic diversity across samples at different MSI loci. Our approach complements existing short tandem repeat callers by preserving read-level diversity and delivering targeted, quantitative MSI calls with potential applications in mechanistic research and clinical assay development.

## Introduction

Microsatellites are highly polymorphic repetitive DNA sequences that exhibit elevated mutation rates ^1^. Their tandem repeat structure renders them particularly prone to insertions and deletions during DNA replication, leading to microsatellite instability (MSI), a hallmark of mismatch repair (MMR) deficiency ^2,3^. Conventional MSI characterization relies on PCR amplification followed by electrophoretic sizing. However, this method provides only average amplicon length estimates from bulk DNA, lacking per-base resolution to identify where and what indels exist, and offers no sequence-level information. Traditional short-read sequencing also faces limitations in accurately resolving MSI regions due to their repetitive nature, resulting in insufficient coverage and limited genomic context.

Recent advances in long-read sequencing technologies have opened new opportunities for high-resolution MSI detection and characterization. Oxford Nanopore Technologies (ONT) sequencing has demonstrated utility in diverse applications, including telomere length and DNA modification profiling ^4–6^. However, ONT’s relatively high error rate (∼5–15%) complicates low-coverage discovery, and most existing error-correction tools are optimized for genome assembly, producing a consensus sequence that collapses intra-sample heterogeneity that may be biologically relevant ^7^. While recent efforts in tandem-repeat genotyping provide robust frameworks for population-level analysis, they lack the depth and resolution required to dissect intra-sample MSI variability ^8,9^. Thus, existing tools have yet to fully leverage deep targeted sequencing to explore MSI heterogeneity.

To address this gap, we present MSIanalyzer, a targeted MSI assay that utilizes ONT to capture and characterize MSI variation at the level of individual sequencing reads. Compared to existing short tandem repeat (STR) callers, our approach provides per-read microsatellite context, applies cluster-aware statistical models, and generates interpretable indel visualizations, enabling quantitative, single-nucleotide resolution analysis of MSI complexity that underlies tumor evolution and susceptibility to diseases associated with repeat instability.

## Methods

### Sample collection and culture methods

We analyzed six samples, including two peripheral blood mononuclear cell (PBMC) samples from anonymous healthy donors, a lymphoblast cell line - TK6, an osteosarcoma cell line - U2OS, two colorectal cancer (CRC) cell lines with known MSI caused by mismatch repair (MMR) deficiencies - HCT15 (MSH6-deficient) and HCT116 (MLH1-deficient). U2OS cells were purchased from ATCC, HCT15 and HCT116 were purchased from the DCDT tumor repository (https://dtp.cancer.gov/repositories/dctdtumorrepository/default.htm) at the National Cancer Institute. TK6 cells were purchased from the TK6 Consortium (http://www.nihs.go.jp/dgm/tk6.html). PBMCs were purchased from a commercial vendor (SMEBio). U2OS and HCT116 cells were cultured in DMEM high glucose with pyruvate (ThermoFisher Cat. No. 11995065) with 10% fetal bovine serum (FBS, ThermoFisher Cat. No. 10437-028). HCT15 was cultured in RPMI 1640 (Gibco, Cat No. 11875085) with 10% FBS. DNA was isolated directly from quiescent PBMC immediately after recovery from cryopreservation.

### Sample preparation and sequencing

Genomic DNA was isolated from approximately at least 1 million cells per sample using QIAGEN DNeasy Blood & Tissue Kit (Qiagen). Quantity and purity were evaluated by spectrophotometry using a Nanodrop One (ThermoFisher).

We targeted five MSI markers from the NCI Bethesda panel: BAT25, BAT26, D2S123, D5S346, and D17S250 ^10^. PCR primer sequences for each marker were obtained from published sources and are listed in **Table 1**. Primer specificity was initially assessed using BLAST against the hg38 reference genome ^11^. The exact genomic positions of these markers were further confirmed via UCSC Genome Browser using the STS Marker track for D2S123, D5S346, and D17S250 and the RepeatMasker track for BAT25 and BAT26 ^12^.

**Table 1.**
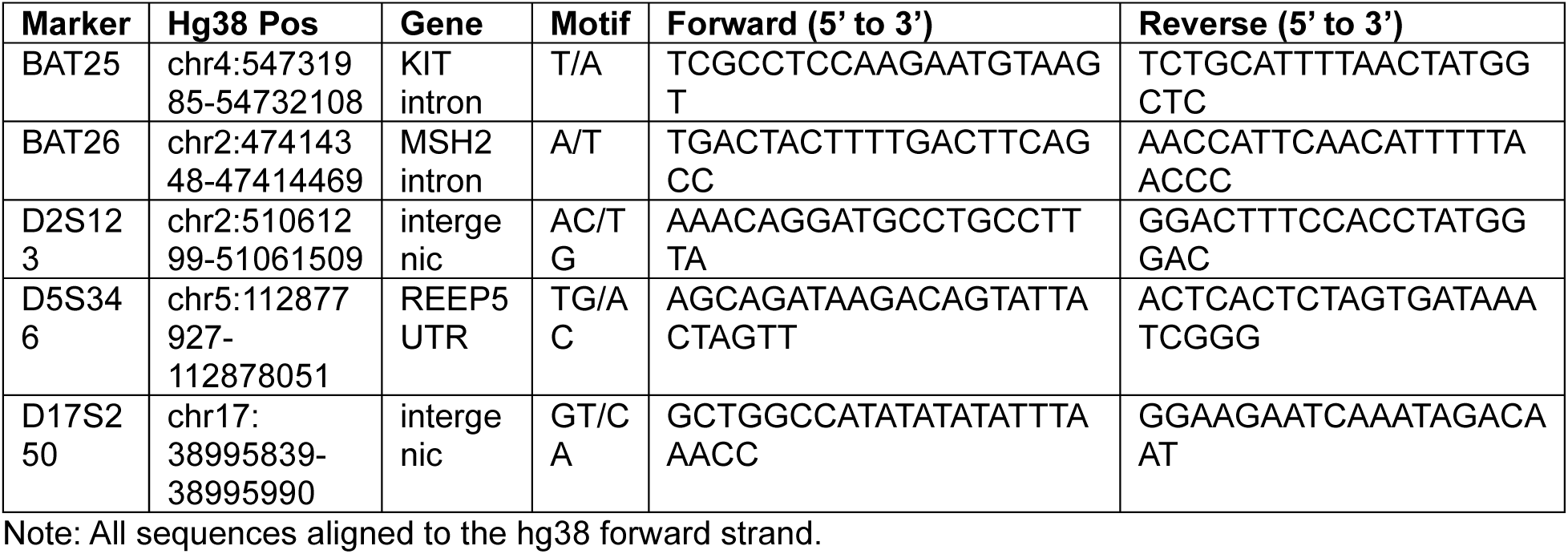
MSI marker details.

PCR was performed using 100 ng of input genomic DNA in a 25 μL total reaction volume, consisting of 12.5 µL NEB 2X Q5 Master Mix, 1.25 μL of each 10 µM primer, 1 μL DNA, and 9 μL nuclease-free water. Thermocycling conditions were: initial denaturation at 98°C for 30 s, followed by 35 cycles of 98°C for 10 s, 60°C for 30 s, and 72°C for 90 s.

PCR products were purified by Monarch PCR Cleanup kit (New England Biolabs), DNA concentration was determined by Nanodrop, and Nanopore sequencing was performed by Plasmidsaurus using the “Premium PCR” option. Samples were prepared according to the vendor’s protocol for amplicon-based Nanopore sequencing, using input concentrations of 20–200 ng/μL in 10 μL. An average of 5,000 reads per sample was obtained.

### Read filtering and primer matching

Raw FASTQ files underwent initial quality control using FastQC (v0.12.1) to verify the presence of targeted marker sequences and assess overall sequencing quality, including Phred score distributions, base composition, and potential adapter contamination. Reads were subsequently filtered to retain only on-target amplification products. To tolerate errors in Nanopore data, we used a Smith-Waterman local alignment with a similarity threshold (sim ≥ 0.85)^13^, which was calculated below with the Levenshtein distance:

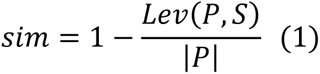

Where *P* is the primer sequence and *S* the sequence segment being aligned. A read is accepted when both primers align and appear in the correct 5’ to 3’ orientation. Because Nanopore reads may be sequenced from either strand, reads aligning to the reverse strand were reoriented to match the forward-strand convention of the hg38 reference genome. Post-filtering yielded >88% of raw input, with median quality scores around Q30, indicating high-quality, on-target amplification.

### Repeat detection and motif counting

To detect microsatellite repeat units, we implemented a robust anchor-extension algorithm customized for Nanopore data. This approach improves upon traditional longest contiguous repeat counting by accommodating typical Nanopore errors such as substitutions and small indels. The process involved the following steps:

**1) Anchor discovery**: Reads were scanned using a sliding window of size k, the motif length. Windows containing ≥ 3 consecutive, perfect repeat units were selected as candidate anchors, following established thresholds in STR detection literature ^14,15^.
**2) Bidirectional extension**: Each anchor was extended in both directions, unit by unit, allowing up to 1 substitution or ≤ 2-unit indels per step. Extensions terminated once these error thresholds were exceeded.
**3) Interruption tracking**: Imperfect repeat units encountered during extension were classified as interruptions. For each interruption, we recorded the relative position, type (substitution, insertion, or deletion), and local sequence context. Each read’s output included repeat region boundaries, total pure repeat units, and a catalog of interruptions.
**4) Variant signature assignment**: Reads containing repeat regions were annotated with variant signatures that summarized both length changes and internal interruptions. Signature categories included: “Perfect[N]”: perfect repeat of *N* units; “Int[N]”: same length with reference but with ≥ 1 internal interruption; “Exp+X” or “Con–X”: expansion or contraction by *X* units relative to the reference. Optional tags denoting interruption types and positions were included if observed at a frequency above a default confidence cutoff of 1%, based on expected Nanopore error rates. Reference allele lengths were defined from a matched reference sample or hg38 annotations.

### Statistical analysis

To ensure robust quantification, additional filters were applied to exclude low-support loci (repeat-region read count <5; flanking-region read count <10), removing ∼10% of the post-filtered reads. Thresholds were selected based on empirical performance and prior studies ^16^.

Our primary objective was to compare repeat unit distributions between samples to identify insertions or deletions at the tandem repeat region. To account for within-sample read correlations, we employed a cluster-aware Dirichlet-multinomial test, which models overdispersed read counts across motif count categories ^17^. Specifically, let 𝑋_𝑟s_ = (𝑥_𝑟s1_, …, 𝑥_𝑟s*K*_) be read-level motif count vector and thus *p* be the proportion of motif count for cluster *r* in sample *s*, for each motif *k* we computed:

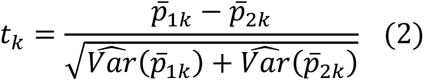

Variances were estimated under a Dirichlet-multinomial model, with the concentration parameter γ̄ derived via maximum likelihood. Multiple testing correction was performed using false discovery rate (FDR) adjustment.

To assess read-level indels, we further classified each read as an indel if its repeat unit count deviated from the modal allele(s) observed in a reference sample. For loci with bi- or multi-modal distributions, defined as any repeat count supported by at least 10% total reads post-filtering, the full set of modal alleles was used to define the reference, and any count outside this set was considered an indel. Indel occurrence rates were modelled with a beta-binomial test to account for overdispersion. The underlying indel proportion in each group was modeled as:

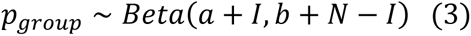

Where *I* is the number of indel-supporting reads, *N* is the total read count, and *a* = *b* = 1 for a non-informative *Beta(1,1)* prior. The posterior distribution of the group-wise difference in indel proportions 𝛥 = 𝑝_1_ − 𝑝_2_ was used as the test statistic. Posterior p-values were estimated as the proportion of sampled posterior draws with Δ>0 or Δ<0.

Additionally, to assess differences in repeat region or amplicon length distributions, we performed non-parametric bootstrap resampling to estimate confidence intervals for length shifts between samples.

### Visualization

Primary visualizations included bar plots of read distributions across repeat units and heatmaps of exact counts across repeat units. To highlight sample-level shifts, we visualized Δ (delta) in motif count proportions between samples. Rare motif lengths in the distribution tails were truncated for visualization but retained in the raw output.

To contextualize variant patterns, genome-aligned pileup plots were generated using trimmed hg38 reference sequences (restricted to PCR target regions) and aligned with Minimap2 (v2.28) in ONT mode suitable for Nanopore sequencing data. Pysam (v0.23.3) was used to extract pileup information, including mismatches and indels, for visualization. Representative reads were further visualized by Integrated Genome Viewer (IGV) to provide per-read contexts ^18^.

### Code availability

MSIanalyzer is open access on GitHub through: https://github.com/NagelLabHub/MSIanalyzer. A demo is provided on Goolge Colab (https://colab.research.google.com/drive/13PjP7rVajoGOFAizytyXdfj6Souv2cts?usp=sharing) for quick analysis with input data.

## Results

### Workflow overview

**Fig. 1** presents the end-to-end MSIanalyzer workflow. Complete analysis of six samples per marker takes approximately 5 minutes, with optional multi-threading available for scalable performance on larger datasets.

**Fig. 1.**
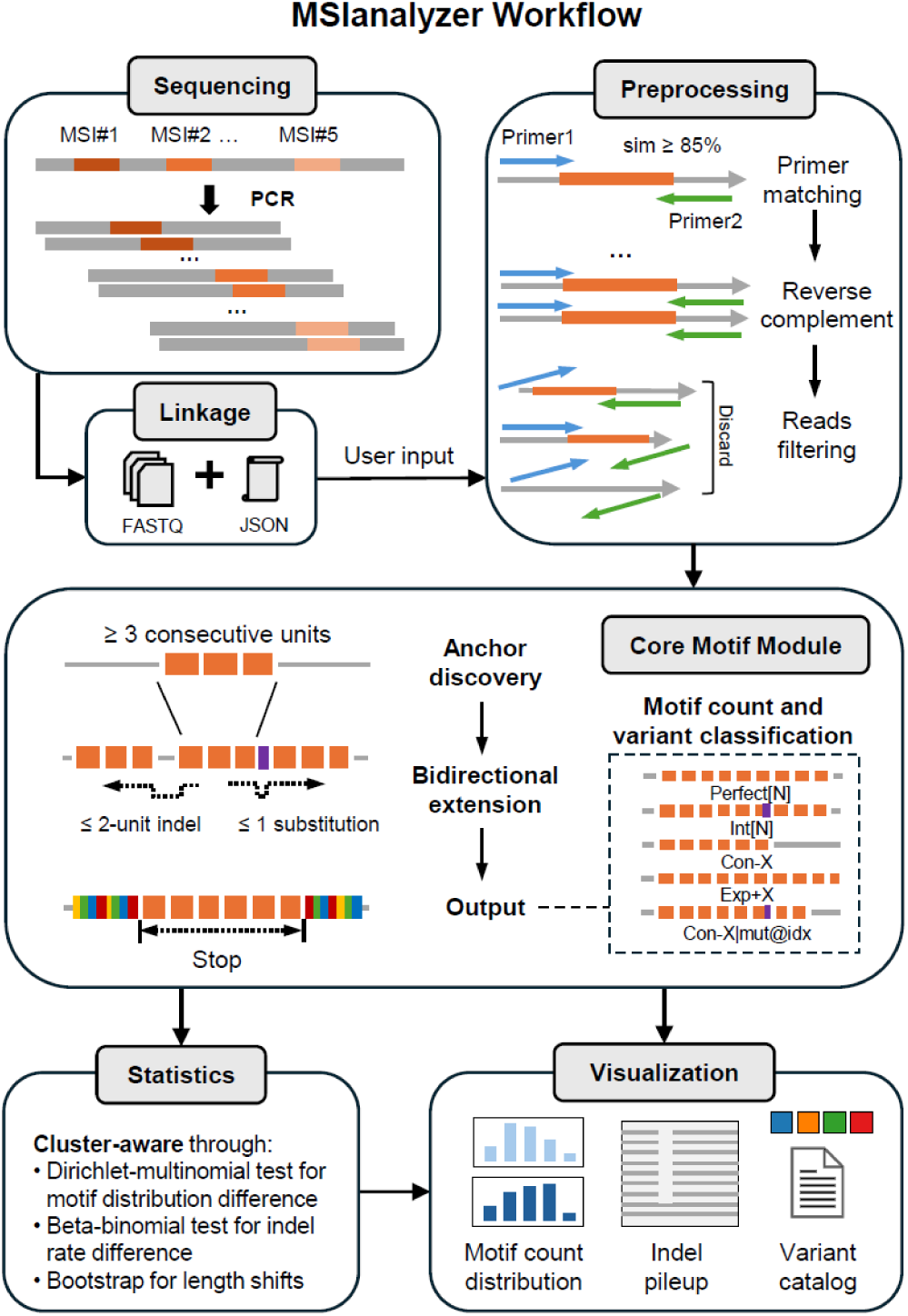
Overview of the MSIanalyzer workflow. Targeted PCR amplicons from microsatellite (MSI) loci are sequenced using Oxford Nanopore Technology (ONT), producing per-sample FASTQ files (Sequencing). A user-supplied JSON manifest links target loci to sequencing data (Linkage). After quality control, reads are matched to their forward and reverse primers with Smith-Waterman alignment (similarity [sim] ≥ 85%), re-oriented to the 5’ to 3’ strand, and off-target reads are discarded (Preprocessing). The core functionality is a robust anchor-extension algorithm, which first detects an anchor of ≥ 3 perfect repeat units, then extends bidirectionally one unit at a time while tolerating ≤ 1 substitution or ≤ 2 indel units per step. Imperfect units encountered during extension are logged in an interruption catalogue (position, edit type, sequence). Final repeat tracts are summarized into variant signatures: Perfect[N], Int[N], Exp+X, or Con−X, with optional annotations (e.g., Con−1 | del@7) if interruptions exceed a defined frequency threshold (see *Methods*). Statistical comparisons account for within-sample read clustering using a Dirichlet-multinomial test (repeat unit distributions), beta-binomial model (indel rates), and bootstrap resampling (length shifts). Visual outputs include bar plots, interruption pileups, stacked variant barplots, and summary tables.

### MSI markers

**Table 1** lists the five Bethesda-panel loci with their hg38 coordinates, gene contexts, repeat motifs, and primer sequences. BAT25 and BAT26 are A/T mononucleotide tracts, whereas D2S123, D5S346 and D17S250 are AC/TG dinucleotide repeats.

### Repeat unit distribution across cell lines

We compared repeat unit profiles among the two CRC cell lines with known MMR defects and four MMR-proficient controls (two cancer cell lines and two PBMC samples). **Fig. 2 and Supplementary Fig. 1** demonstrate clear modal shifts separating MMR-deficient from MMR-intact samples, with p < 0.05 from beta-binomial tests of indel rates compared to the PBMC reference across the five markers (**Supplementary Table 1**). Most loci displayed high read frequency at multiple repeat units, with 2 to 4 repeat unit lengths supported by at least 10% of reads (**Supplementary Tables 2-6**). PBMC samples also exhibited allelic variation among themselves, with differential indel rates detected by beta-binomial tests at the dinucleotide loci.

**Fig. 2.**
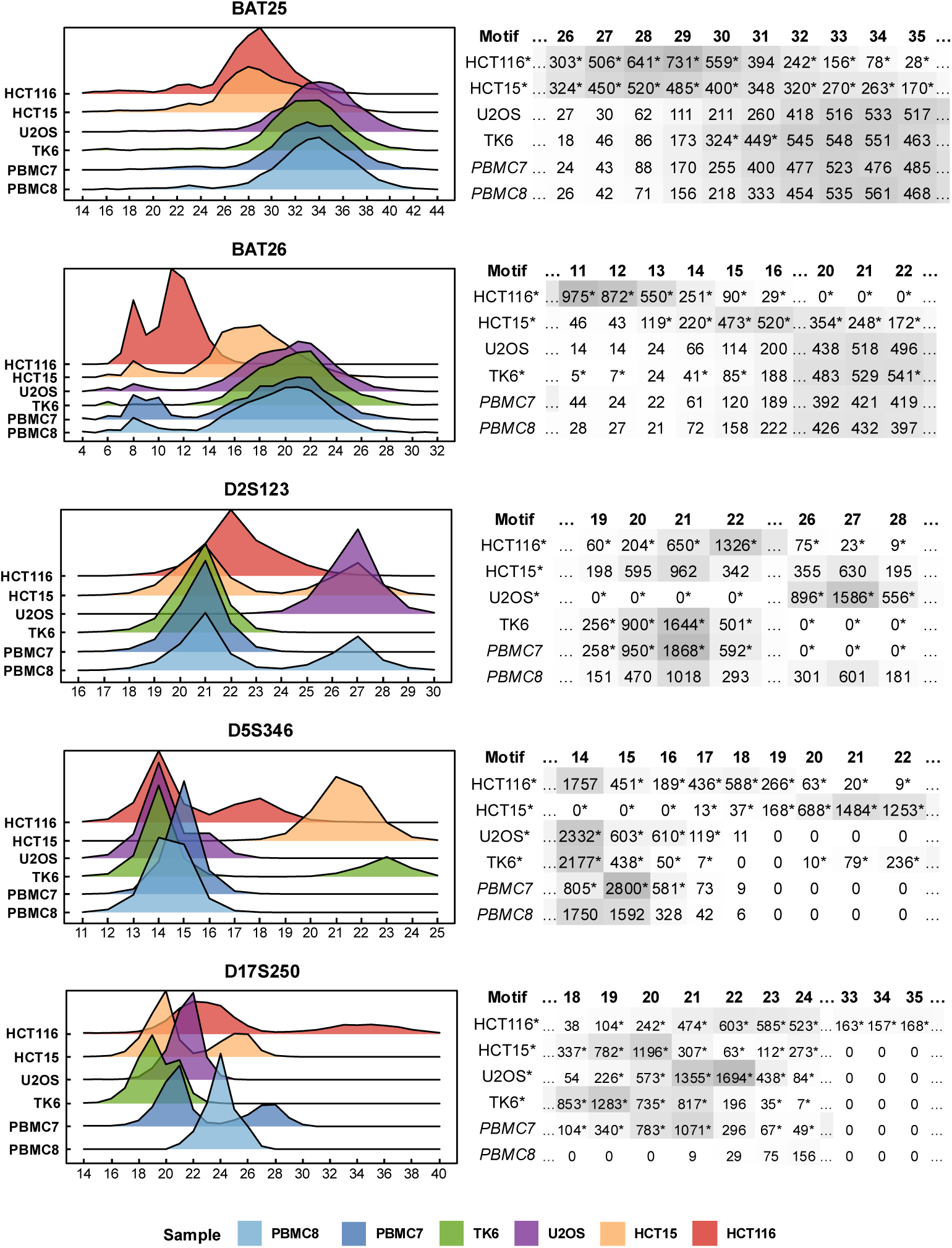
Repeat unit distribution across samples. **Left panels**: Smoothed ridge plots showing normalized read frequencies at each repeat unit count for each sample (color-coded). Curves highlight modal shifts and distribution breadth across markers. **Right panels**: Heatmaps of raw read counts (darker gray = higher counts) across repeat unit bins and samples. Sparse tails are truncated to reduce clutter. Asterisks (*) inside the cells indicate significance (FDR < 0.05) by Dirichlet-multinomial test vs. PBMC8. Asterisks next to sample names indicate a significant indel rate difference (p < 0.05, beta-binomial test) compared to both PBMC7 and PBMC8.

At the mononucleotide (T/A) repeat **BAT25**, both HCT15 and HCT116 contracted to modes at 28–29 repeat units, with mean reductions of 4 units for HCT15 (95% CI: –4.24 to –3.91) and 5 units for HCT116 (95% CI: –5.33 to –5.01) relative to PBMCs. Dirichlet-multinomial tests confirmed significant under-representation of units 32–35 in both samples (FDR < 0.05), whereas TK6 and U2OS profiles overlapped those of PBMCs.

At the mononucleotide (A/T) repeat **BAT26**, HCT116 showed a pronounced contraction tail centered at 11–13 units (FDR < 0.05), yielding a mean shift of –8 units (95%CI: –8.13, –7.823). HCT15 contracted more modestly to 15–16 units (mean: −2.1 units, 95% CI: –2.26, –1.90; FDR < 0.05). TK6 and U2OS aligned closely with the PBMC profile. A minor peak at 8 units was detectable in PBMCs and likely represents a rare polymorphism described in prior studies ^19^. Its higher frequency in HCT116 suggests instability contributing to contraction to this length.

For the dinucleotide (AC/TG) repeat **D2S123**, several lines displayed bimodal distributions with peaks at 21 and 27 units. U2OS was almost exclusively centered at 27 units (FDR < 0.05 for units 26–28), whereas HCT116 retained a unimodal 22-unit profile that differed from the modal 21-unit profile in PBMCs (FDR < 0.05 at 21–22 units). HCT15 mirrored the PBMC8 unimodal profile, while TK6 resembled the PBMC7 bimodal profile.

At the dinucleotide (TG/AC) repeat **D5S346**, HCT15 alone showed a clear expansion peak at 21–22 units, a gain of 6.9 units (95%CI: 6.819, 6.907, FDR < 0.05) compared to PBMC. Other lines retained the 14–16 unit range, although HCT116 showed a secondary peak at 17–18 units (FDR < 0.05).

The dinucleotide (GT/CA) repeat **D17S250** exhibited the greatest heterogeneity: HCT116 reads spanned 19–40 units, with 14 individual repeat units showing significant differences when compared to PBMC8 (FDR < 0.05), consistent with previous observations of extensive instability at this locus ^20^. Modal diversity was also observed across the remaining samples, with PBMC samples being bimodal at 20 and 24 units, TK6 peaked at 19, U2OS at 22, and HCT15 at 20 with a shoulder at 25–26.

### Per-read variability in indels distribution across cell lines

Indel and mismatch rates by genomic position are summarized in **Supplementary** Fig. 3-7. To visualize per-read variability, we examined BAT26 read stacks in IGV. **Fig. 3** contrasts TK6 and HCT116: in TK6, reads showed mostly uniform deletions <10 bp relative to hg38. HCT116 exhibited greater variability, with deletion lengths ranging from 12–18 bp. This further illustrates MSIanalyzer’s ability to capture per-read interruption heterogeneity at single-molecule resolution.

**Fig. 3.**
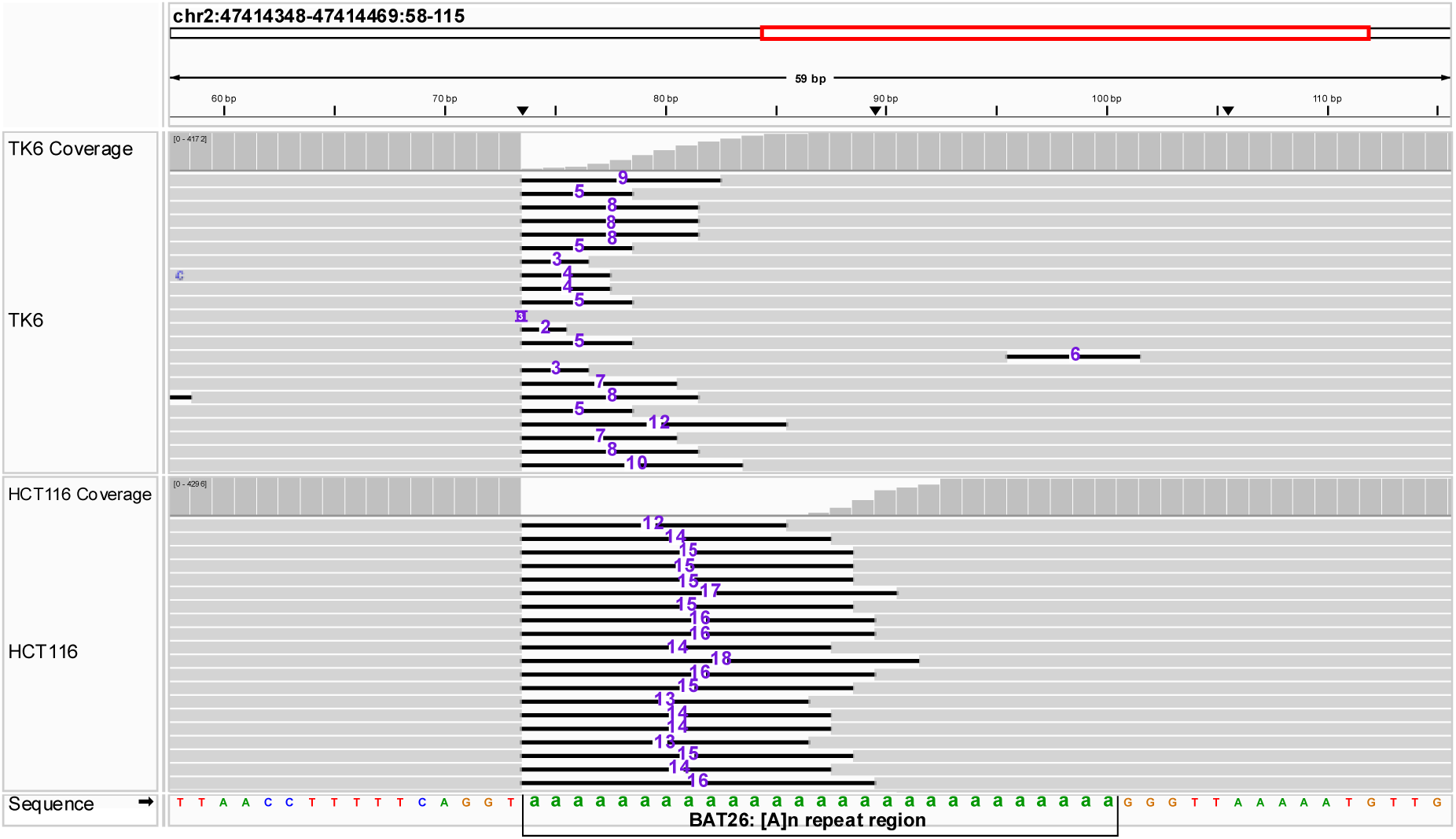
Read-level indel diversity at BAT26. IGV snapshots show read alignments at BAT26 for TK6 and HCT116. Top: trimmed hg38 reference. Grey: individual reads. Black: deletions in individual reads. Purple numerals: deletion lengths (bp) relative to hg38. Green bases denote the targeted mononucleotide region.

## Discussion

Our results demonstrate that MSIanalyzer accurately captures hallmark MSI patterns in MMR-deficient cell lines and reveals locus-specific differences between MLH1- and MSH6-deficient backgrounds and MMR-proficient controls. Operating directly on sequencing reads, the pipeline preserves per-molecule information, accounts for platform-specific errors without collapsing underlying biological heterogeneity, and evaluates between-sample differences using statistical models that accommodate read clustering. In addition, the ability to visualize read-level interruption patterns enables direct quantification of indel length and frequency. These features make MSIanalyzer a broadly applicable platform not only for MSI profiling but also for other STRs.

Several existing tools have advanced the detection of STR variation across platforms. NanoRepeat effectively identifies trinucleotide expansions but is less suited to interruption-rich loci and relies on alignment ^21^. Tandem Repeats Finder is widely used for initial STR screening but does not support interruption-aware or quantitative comparisons ^22^. NanoSTR is specifically designed for ONT data but lacks per-read diversity reporting ^23^. OS-Seq enables high-accuracy, multiplex MSI profiling using custom probe-based capture and primer extension, though it relies on specialized oligonucleotide synthesis and instrumentation, which may limit flexibility for broader applications ^24^. MSIanalyzer complements these tools by focusing on preserving intra-sample allelic variation and read-level variation, enabling single-molecule resolution and visualization of MSI patterns.

We acknowledge several limitations of our approach. First, as a targeted approach relying on PCR amplification, MSIanalyzer is susceptible to amplification bias and polymerase slippage, potentially distorting allele spectra, particularly at very long homopolymers. Second, this proof-of-concept study includes only six samples, limiting power to detect rare alleles. However, the implementation of a Dirichlet-multinomial test helps address overdispersion. Third, we observed discrepancies in motif counts compared to the hg38 reference genome, likely stemming from persistent platform-specific sequencing artifacts. Without external validation or larger sample cohorts, these cannot be definitively resolved. Nonetheless, within-sample comparisons remain robust, as all samples are subject to the same artifacts, which are mitigated through a robust analytical framework.

Despite these limitations, our results reveal locus-specific distinctions between MLH1- and MSH6-deficient CRC cell lines and MMR-proficient controls, supporting that different DNA repair deficiencies produce characteristic MSI signatures ^25^. If confirmed in larger cohorts, these patterns could inform tumor classification and enhance biomarker discovery for precision oncology. Future development of MSIanalyzer will focus on benchmarking against complex repeat regions in population-scale datasets and validate the statistical framework under real-world heterogeneity. With continued refinement, MSIanalyzer has the potential to become a general-purpose, low-cost solution for research and clinical MSI typing.

## Acknowledgments

This research was partially supported by a sponsored research agreement with Pfizer Inc. ZDN and DJL also acknowledge support from American Cancer Society grant (RSG-22-038-01) and NIH grant (P01CA092584).

## Declaration of Interests

Z.D.N. received funding for this project from Pfizer Inc., holds a patent titled “Methods and kits for determining DNA repair capacity,” and has received research funding from Intellia Therapeutics, Ensoma, and Agios that is unrelated to this project.

## Author Contributions

Conceptualization and methodology: TZ, DJL, ZDN

Investigation: TZ, DJL

Software, formal analysis, visualization, writing – original draft: TZ

Resources, writing – review and revision: DJL, ZDN

Supervision and funding acquisition: ZDN

---

**Supplementary Fig. 1.**
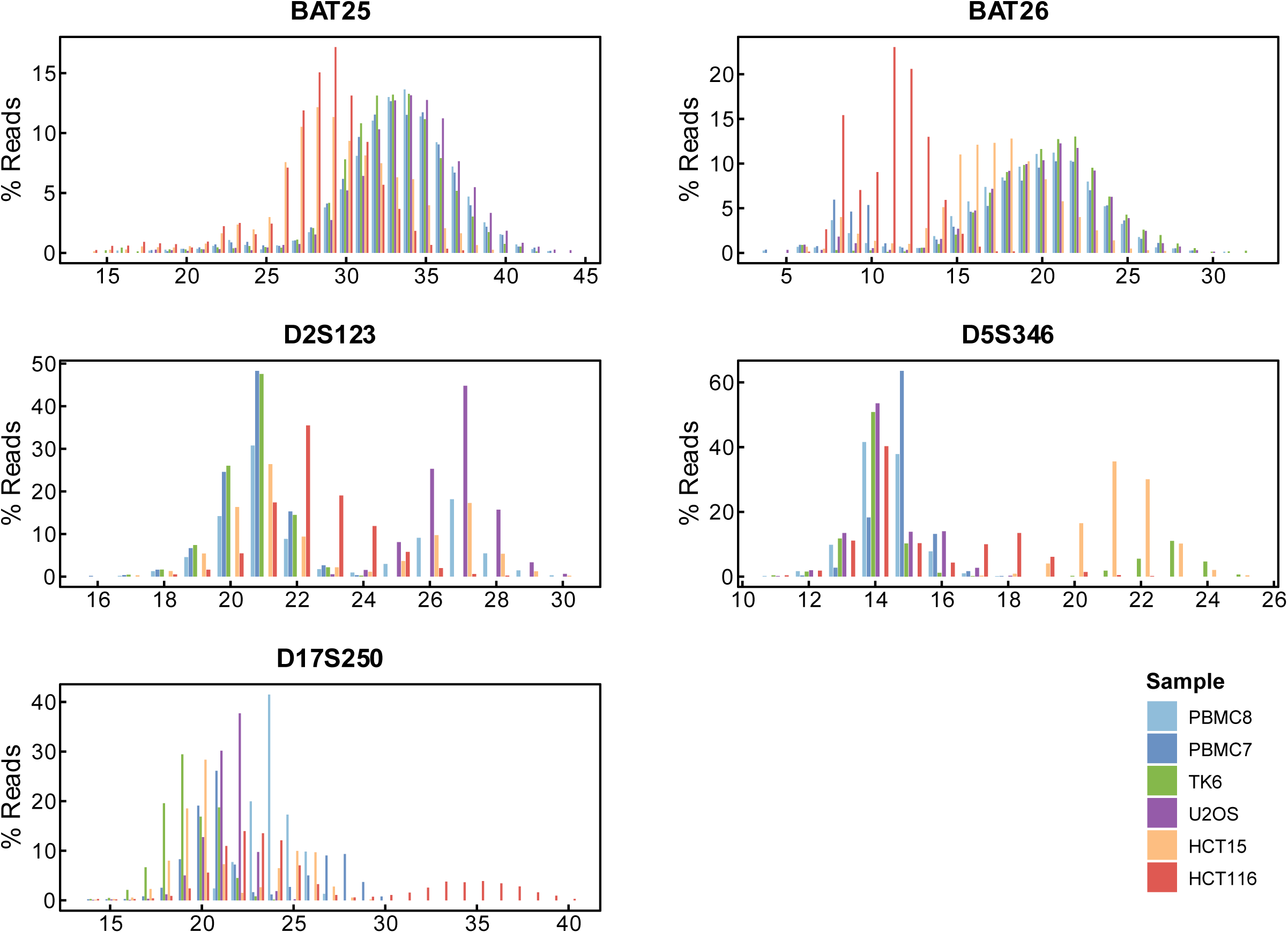
Repeat unit distribution across samples. Each bar represents the percentage of reads observed at a given repeat unit, grouped by sample and color-coded as indicated in the legend.

**Supplementary Fig. 2.**
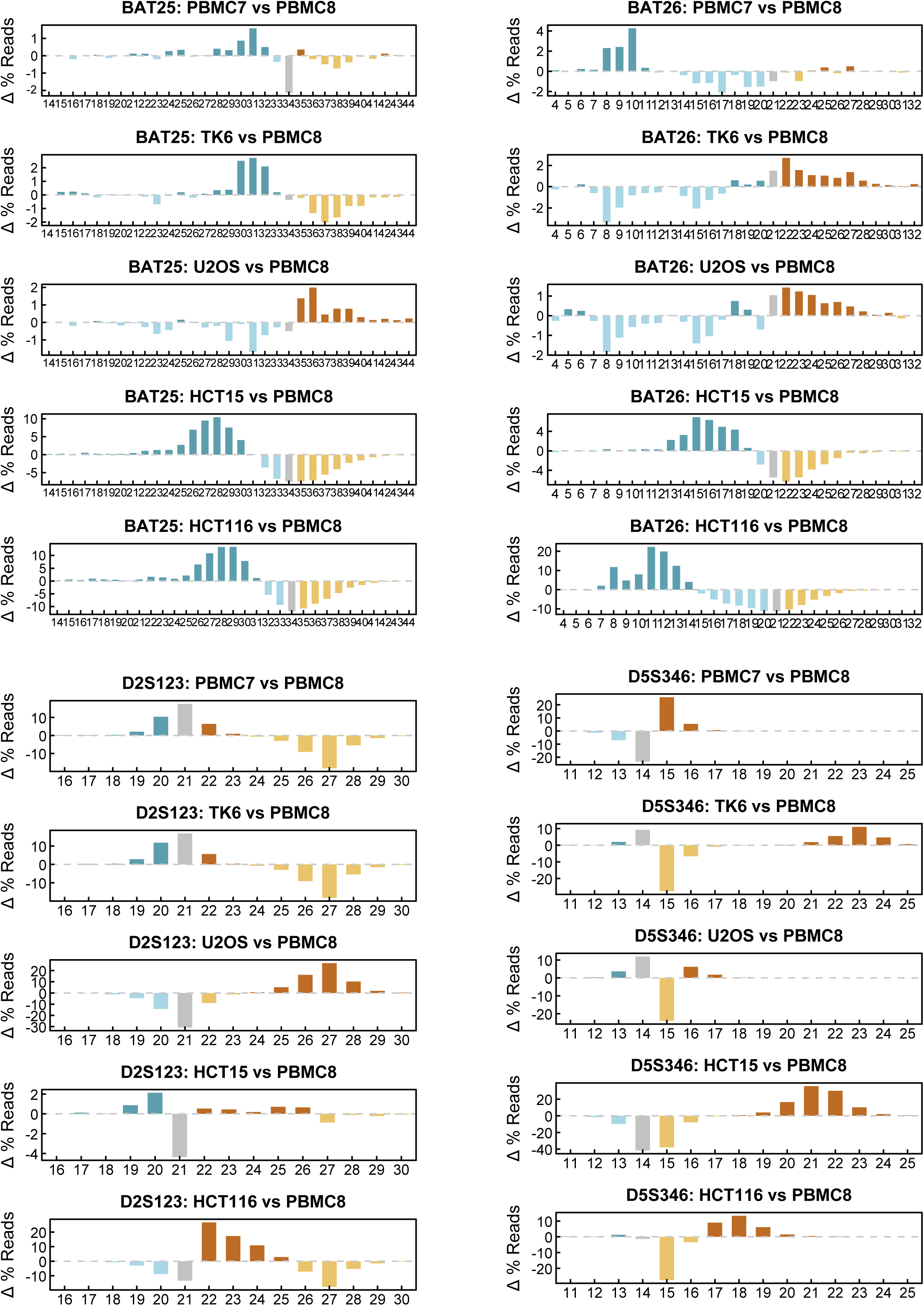
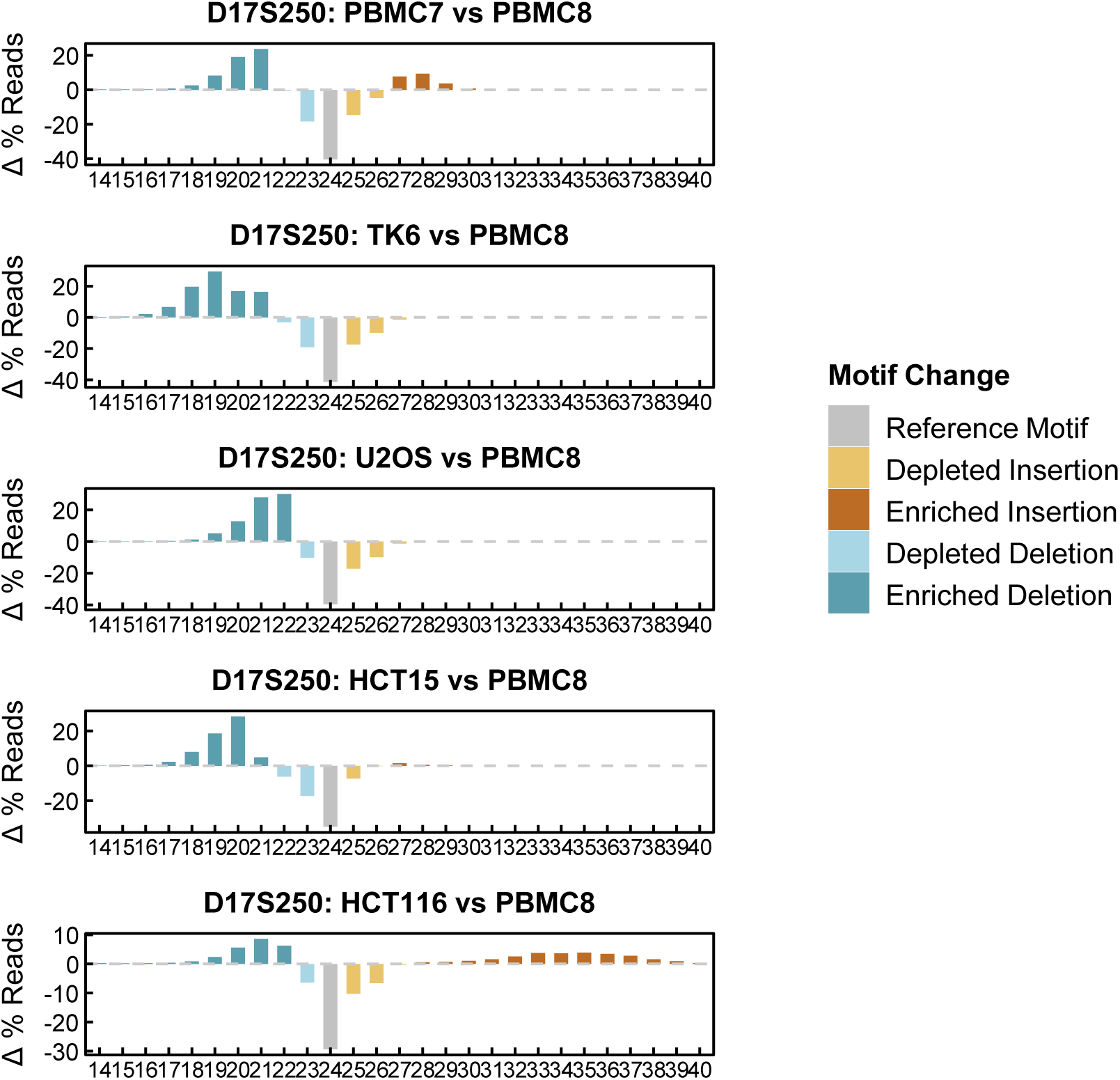
Difference in repeat unit proportions across samples. The difference in read proportion (Δ % Reads) for each sample compared to the PBMC8 reference. Bars show differences in repeat unit frequencies, colored by indel classification: enriched or depleted insertions/deletions, relative to the reference, as defined in the Motif Change key. Gray bars represent reference motif counts (most frequent in PBMC8).

**Supplementary Fig. 3.**
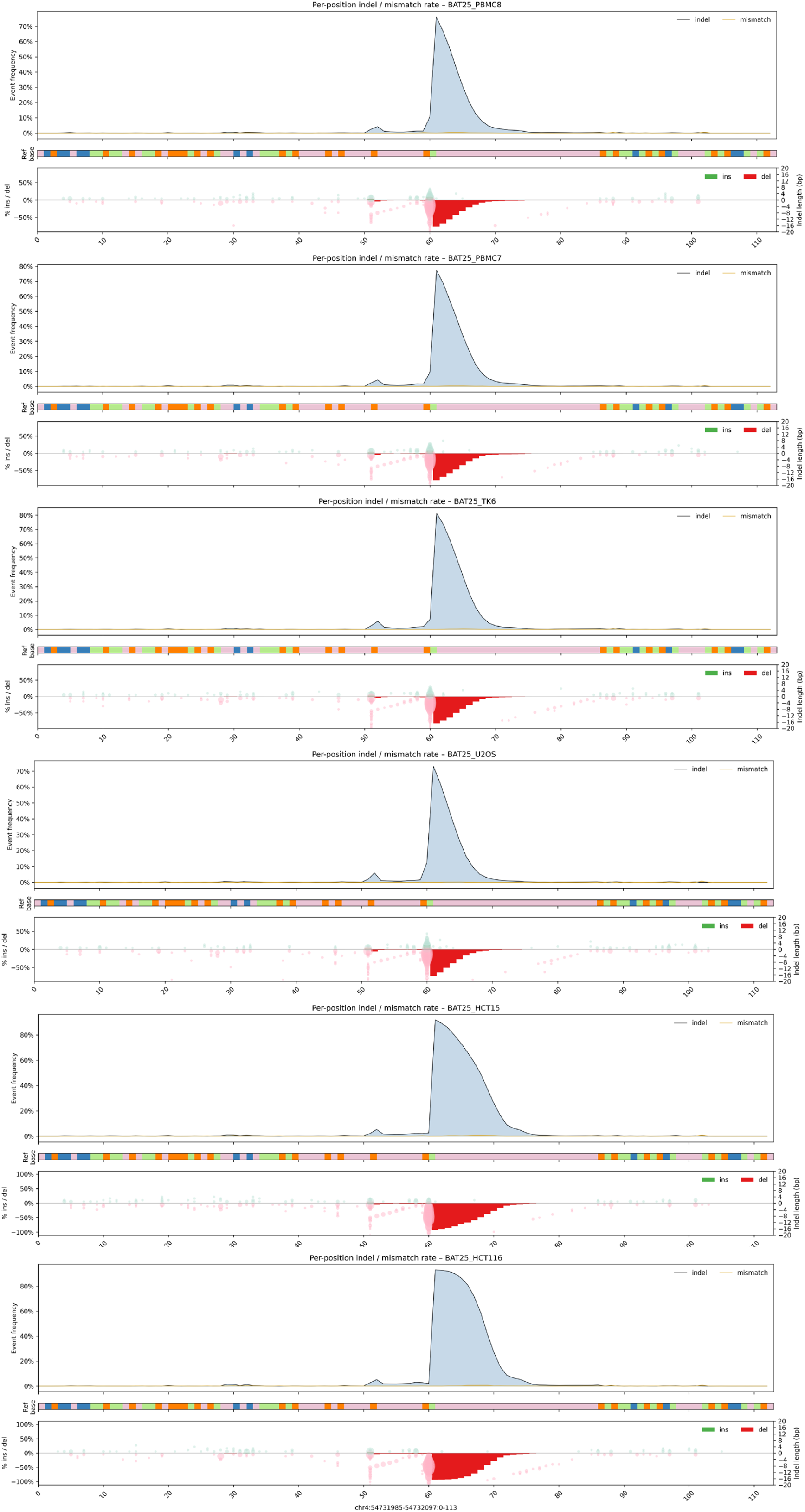
Per-base indel and mismatch pileup profiles at BAT25. **Top panel**: smoothed frequency of indels (blue) and mismatches (gold) along the locus. **Middle panel**: hg38 reference base track. **Bottom panel**: detailed indel visualization showing insertion and deletion types, lengths (in bp), and positions across reads.

**Supplementary Fig. 4.**
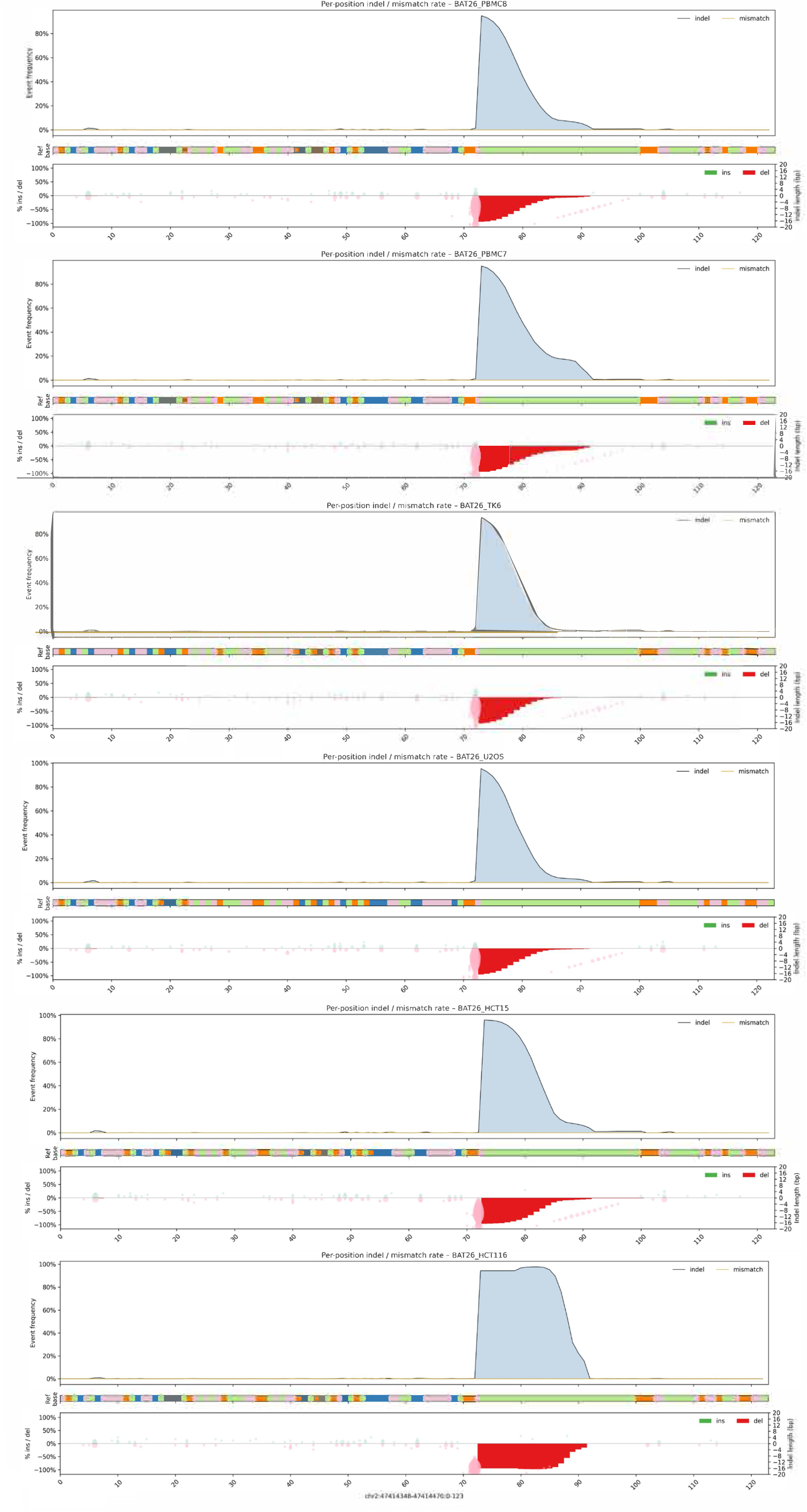
Per-base indel and mismatch pileup profiles at BAT26. **Top panel**: smoothed frequency of indels (blue) and mismatches (gold) along the locus. **Middle panel**: hg38 reference base track. **Bottom panel**: detailed indel visualization showing insertion and deletion types, lengths (in bp), and positions across reads.

**Supplementary Fig. 5.**
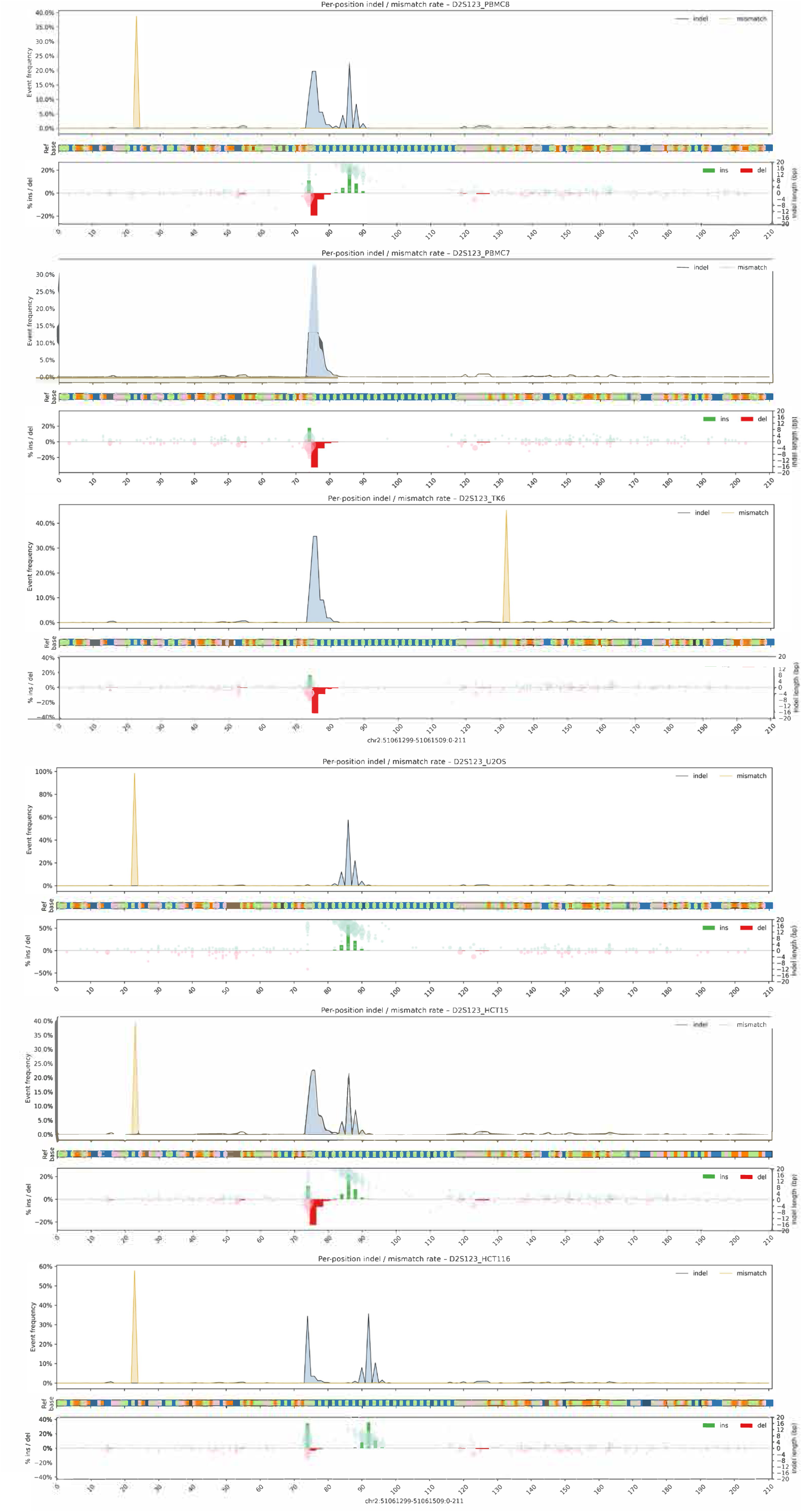
Per-base indel and mismatch pileup profiles at D2S123. **Top panel**: smoothed frequency of indels (blue) and mismatches (gold) along the locus. **Middle panel**: hg38 reference base track. **Bottom panel**: detailed indel visualization showing insertion and deletion types, lengths (in bp), and positions across reads.

**Supplementary Fig. 6.**
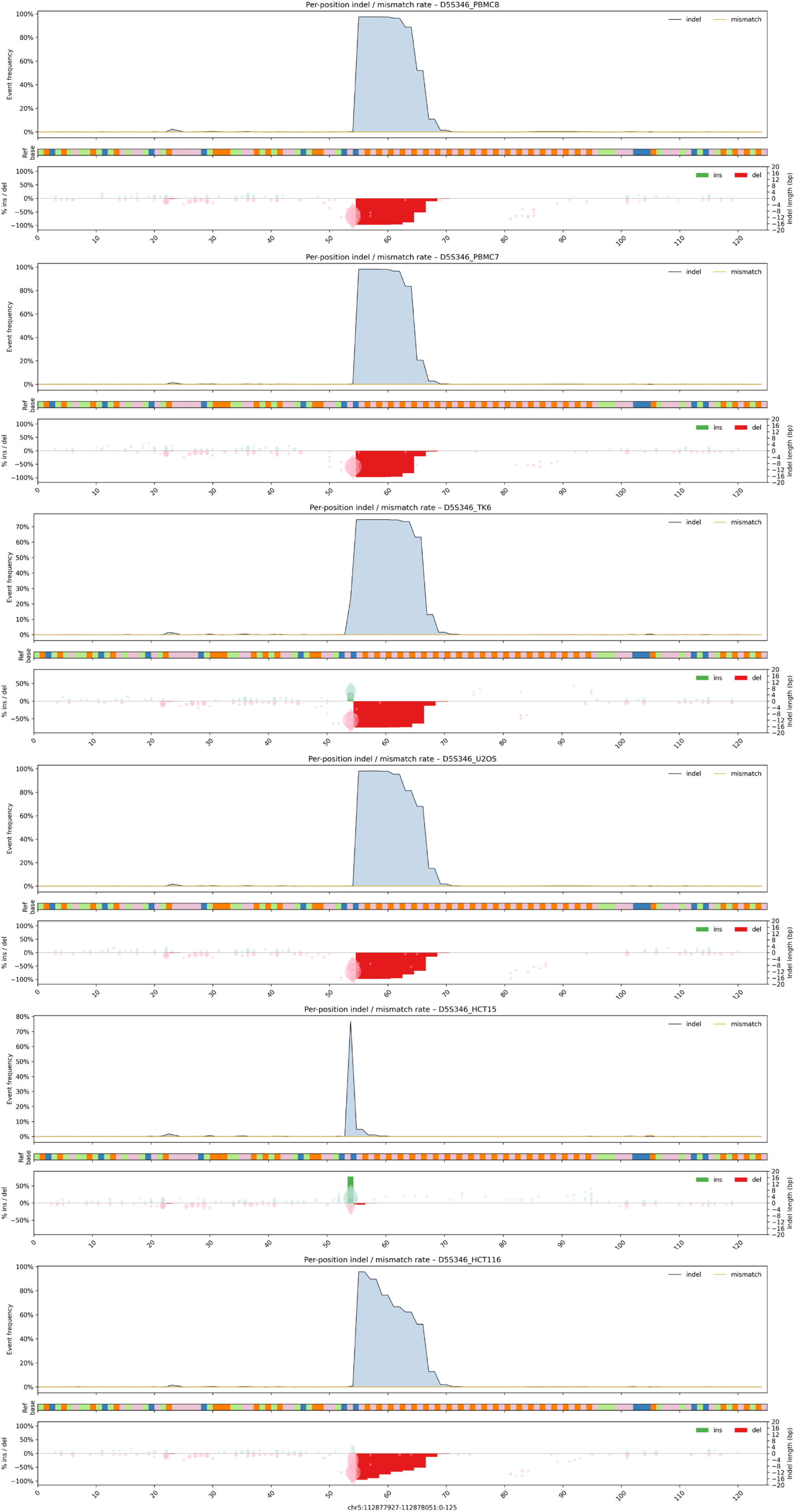
Per-base indel and mismatch pileup profiles at D5S346. **Top panel**: smoothed frequency of indels (blue) and mismatches (gold) along the locus. **Middle panel**: hg38 reference base track. **Bottom panel**: detailed indel visualization showing insertion and deletion types, lengths (in bp), and positions across reads.

**Supplementary Fig. 7.**
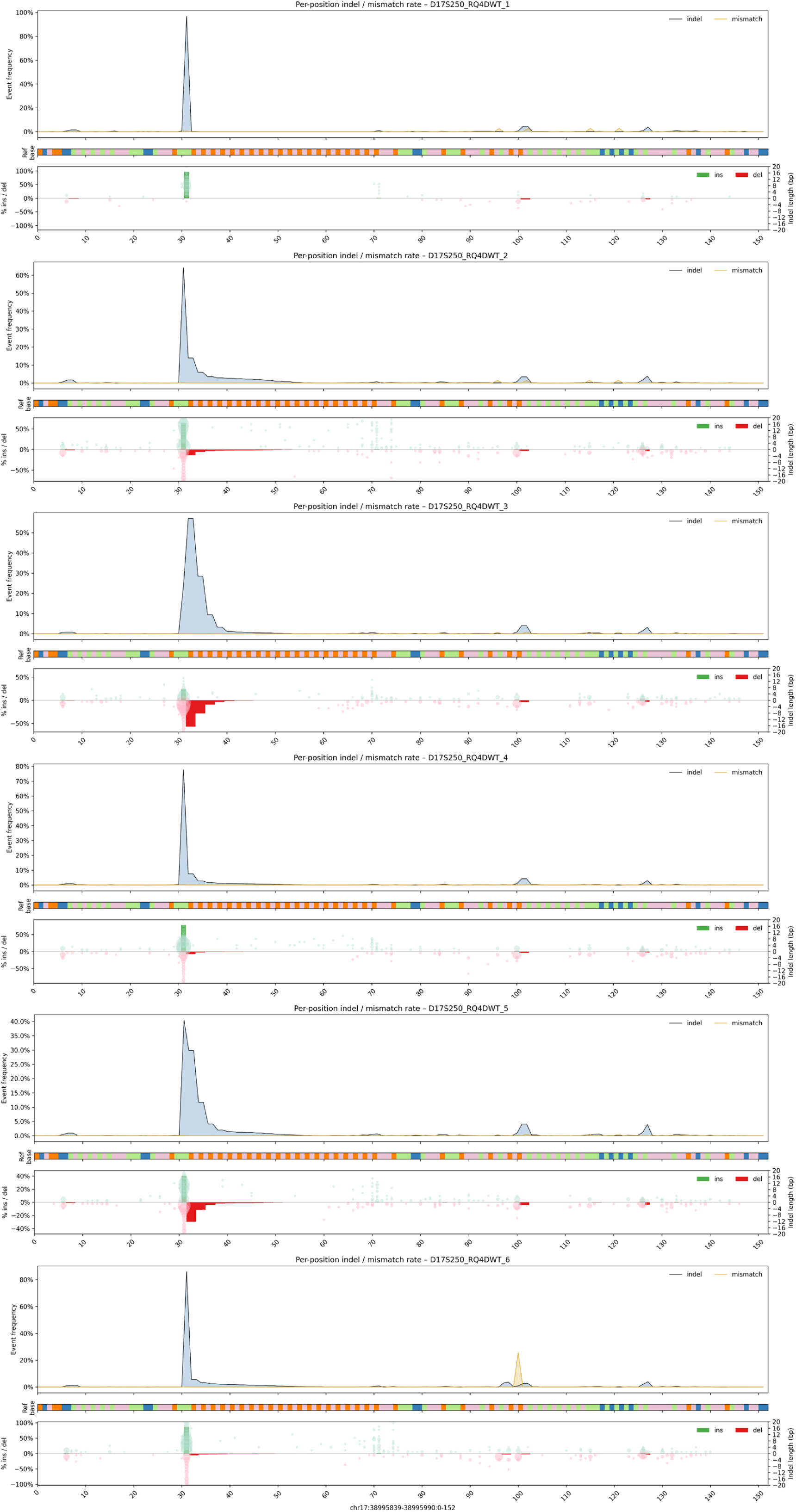
Per-base indel and mismatch pileup profiles at D17S250. **Top panel**: smoothed frequency of indels (blue) and mismatches (gold) along the locus. **Middle panel**: hg38 reference base track. **Bottom panel**: detailed indel visualization showing insertion and deletion types, lengths (in bp), and positions across reads.

**Supplementary Table 1.** Statistical comparison for indel rate and repeat unit length shifts across samples.

|  | Indel rate beta-binomial test |  |  |  | Unit length shift (bootstrapping) |  |  |  |
| --- | --- | --- | --- | --- | --- | --- | --- | --- |
| Reference | PBMC8 |  | PBMC7 |  | PBMC8 |  | PBMC7 |  |
| Sample by marker | rate diff | 95%CI | rate diff | 95%CI | mean | 95%CI | mean | 95%CI |
| BAT25 |  |  |  |  |  |  |  |  |
| HCT116 | 0.37 | (0.354, 0.39) | 0.36 | (0.338, 0.374) | -5.17 | (-5.331, -5.012) | -4.96 | (-5.116, -4.805) |
| HCT15 | 0.25 | (0.231, 0.271) | 0.24 | (0.216, 0.255) | -4.07 | (-4.236, -3.905) | -3.86 | (-4.021, -3.695) |
| U2OS | 0.00 | (-0.02, 0.023) | -0.02 | (-0.037, 0.007) | 0.56 | (0.4, 0.719) | 0.77 | (0.611, 0.928) |
| TK6 | -0.02 | (-0.039, 0.005) | -0.03 | (-0.055, -0.012) | -0.53 | (-0.688, -0.367) | -0.32 | (-0.472, -0.161) |
| PBMC7 | 0.02 | (-0.005, 0.038) | - | - | -0.21 | (-0.372, -0.05) | - | - |
| Canonical alleles | 32, 33, 34, 35 |  | 32, 33, 34, 35 |  |  |  |  |  |
| BAT26 |  |  |  |  |  |  |  |  |
|  | rate diff | 95%CI | rate diff | 95%CI | mean | 95%CI | mean | 95%CI |
| HCT116 | 0.33 | (0.311, 0.341) | 0.20 | (0.192, 0.217) | -7.98 | (-8.13, -7.823) | -7.13 | (-7.305, -6.954) |
| HCT15 | 0.15 | (0.127, 0.165) | 0.11 | (0.091, 0.122) | -2.08 | (-2.263, -1.895) | -1.23 | (-1.431, -1.024) |
| U2OS | -0.02 | (-0.038, 0.003) | -0.04 | (-0.054, -0.018) | 0.82 | (0.634, 1.012) | 1.67 | (1.469, 1.876) |
| TK6 | -0.05 | (-0.068, -0.027) | -0.05 | (-0.071, -0.035) | 1.57 | (1.386, 1.747) | 2.42 | (2.218, 2.617) |
| PBMC7 | 0.03 | (0.006, 0.047) | - | - | -0.85 | (-1.067, -0.632) | - | - |
| Canonical alleles | 20, 21, 22 |  | 21, 22 |  |  |  |  |  |
| D2S123 |  |  |  |  |  |  |  |  |
|  | rate diff | 95%CI | rate diff | 95%CI | mean | 95%CI | mean | 95%CI |
| HCT116 | 0.40 | (0.375, 0.418) | 0.30 | (0.279, 0.317) | -0.69 | (-0.804, -0.572) | 1.61 | (1.553, 1.668) |
| HCT15 | 0.03 | (0.008, 0.054) | 0.36 | (0.341, 0.379) | -0.05 | (-0.194, 0.092) | 2.25 | (2.144, 2.357) |
| U2OS | 0.18 | (0.16, 0.207) | 0.88 | (0.871, 0.891) | 3.70 | (3.586, 3.807) | 6.00 | (5.949, 6.043) |
| TK6 | -0.10 | (-0.127, -0.082) | 0.00 | (-0.014, 0.016) | -2.34 | (-2.455, -2.234) | -0.05 | (-0.092, -0.002) |
| PBMC7 | -0.10 | (-0.119, -0.075) | - | - | -2.30 | (-2.406, -2.188) | - | - |
| Canonical alleles | 20, 21, 27 |  | 20, 21, 22 |  |  |  |  |  |
| D5S346 |  |  |  |  |  |  |  |  |
|  | rate diff | 95%CI | rate diff | 95%CI | mean | 95%CI | mean | 95%CI |
| HCT116 | 0.29 | (0.269, 0.307) | 0.31 | (0.293, 0.331) | 0.88 | (0.81, 0.945) | 0.39 | (0.322, 0.452) |
| HCT15 | 0.79 | (0.781, 0.806) | 0.82 | (0.807, 0.829) | 6.86 | (6.819, 6.907) | 6.37 | (6.334, 6.416) |
| U2OS | 0.12 | (0.102, 0.139) | 0.14 | (0.127, 0.162) | -0.10 | (-0.144, -0.061) | -0.59 | (-0.63, -0.554) |
| TK6 | 0.18 | (0.164, 0.203) | 0.21 | (0.189, 0.226) | 1.65 | (1.532, 1.764) | 1.16 | (1.043, 1.279) |
| PBMC7 | -0.02 | (-0.041, -0.007) | - | - | 0.49 | (0.455, 0.524) | - | - |
| Canonical alleles | 14, 15 |  | 14, 15 |  |  |  |  |  |
| D17S250 |  |  |  |  |  |  |  |  |
|  | rate diff | 95%CI | rate diff | 95%CI | mean | 95%CI | mean | 95%CI |
| HCT116 | 0.46 | (0.414, 0.501) | 0.29 | (0.268, 0.306) | 1.56 | (1.35, 1.769) | 3.18 | (2.969, 3.385) |
| HCT15 | 0.59 | (0.551, 0.637) | 0.10 | (0.074, 0.117) | -2.74 | (-2.89, -2.585) | -1.12 | (-1.271, -0.975) |
| U2OS | 0.67 | (0.624, 0.708) | 0.02 | (0.002, 0.044) | -2.71 | (-2.841, -2.585) | -1.10 | (-1.222, -0.978) |
| TK6 | 0.78 | (0.733, 0.816) | 0.10 | (0.075, 0.117) | -4.72 | (-4.848, -4.594) | -3.11 | (-3.23, -2.982) |
| PBMC7 | 0.73 | (0.687, 0.77) | - | - | -1.61 | (-1.779, -1.448) | - | - |
| Canonical alleles | 23, 24, 25 |  | 20, 21 |  |  |  |  |  |

**Supplementary Table 2.**
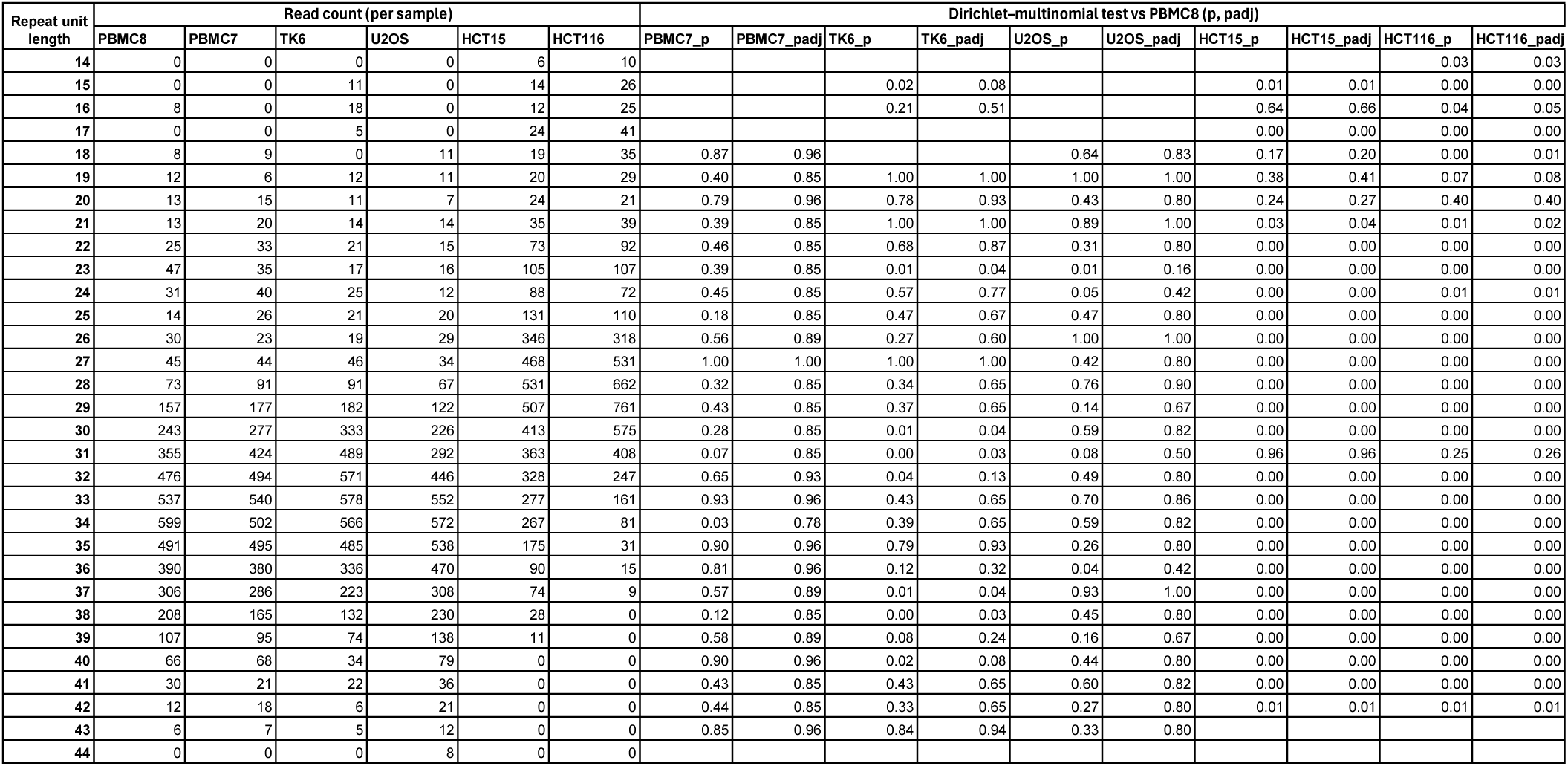
Repeat unit distribution and statistical comparisons for BAT25 across six samples.

**Supplementary Table 3.**
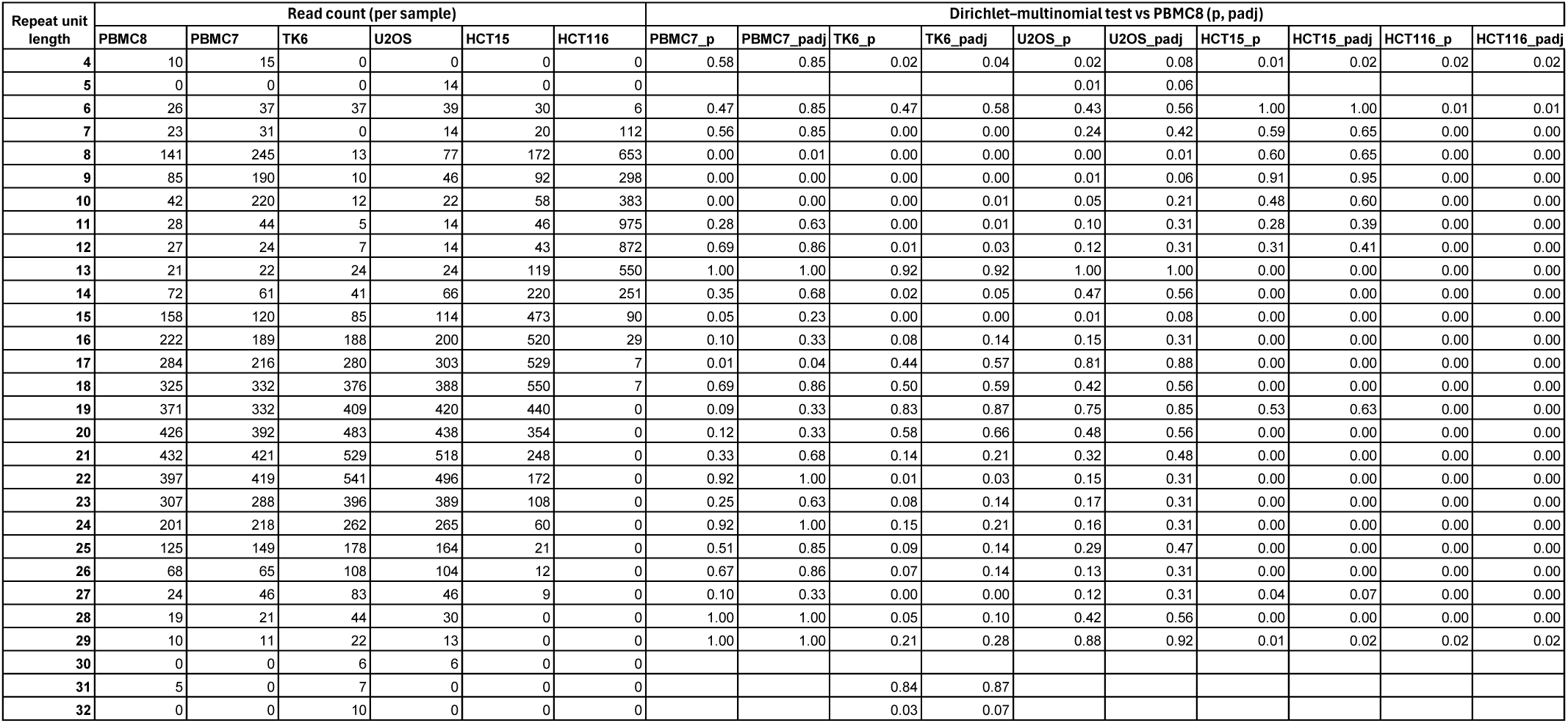
Repeat unit distribution and statistical comparisons for BAT26 across six samples.

**Supplementary Table 4.** Repeat unit distribution and statistical comparisons for D2S123 across six samples.

| Repeat unit length | Read count (per sample) |  |  |  |  |  | Dirichlet-multinomial test vs PBMC8 (p, padj) |  |  |  |  |  |  |  |  |  |
| --- | --- | --- | --- | --- | --- | --- | --- | --- | --- | --- | --- | --- | --- | --- | --- | --- |
|  | PBMC8 | PBMC7 | TK6 | U2OS | HCT15 | HCT116 | PBMC7_p | PBMC7_padj | TK6_p | TK6_padj | U2OS_p | U2OS_padj | HCT15_p | HCT15_padj | HCT116_p | HCT116_padj |
| 16 | 0 | 7 | 0 | 0 | 0 | 0 |  |  |  |  |  |  |  |  |  |  |
| 17 | 6 | 14 | 16 | 0 | 11 | 0 | 0.34 | 0.37 | 0.17 | 0.20 |  |  | 0.50 | 0.79 |  |  |
| 18 | 42 | 63 | 57 | 0 | 47 | 19 | 0.40 | 0.40 | 0.39 | 0.39 | 0.00 | 0.00 | 1.00 | 1.00 | 0.02 | 0.02 |
| 19 | 151 | 258 | 256 | 0 | 198 | 60 | 0.01 | 0.01 | 0.00 | 0.00 | 0.00 | 0.00 | 0.24 | 0.79 | 0.00 | 0.00 |
| 20 | 470 | 950 | 900 | 0 | 595 | 204 | 0.00 | 0.00 | 0.00 | 0.00 | 0.00 | 0.00 | 0.09 | 0.60 | 0.00 | 0.00 |
| 21 | 1018 | 1868 | 1644 | 0 | 962 | 650 | 0.00 | 0.00 | 0.00 | 0.00 | 0.00 | 0.00 | 0.00 | 0.06 | 0.00 | 0.00 |
| 22 | 293 | 592 | 501 | 0 | 342 | 1326 | 0.00 | 0.00 | 0.00 | 0.00 | 0.00 | 0.00 | 0.60 | 0.79 | 0.00 | 0.00 |
| 23 | 58 | 103 | 75 | 19 | 80 | 711 | 0.07 | 0.08 | 0.39 | 0.39 | 0.00 | 0.00 | 0.36 | 0.79 | 0.00 | 0.00 |
| 24 | 32 | 13 | 8 | 55 | 42 | 443 | 0.02 | 0.02 | 0.01 | 0.01 | 0.13 | 0.14 | 0.62 | 0.79 | 0.00 | 0.00 |
| 25 | 98 | 0 | 0 | 287 | 134 | 217 | 0.00 | 0.00 | 0.00 | 0.00 | 0.00 | 0.00 | 0.26 | 0.79 | 0.00 | 0.00 |
| 26 | 301 | 0 | 0 | 896 | 355 | 75 | 0.00 | 0.00 | 0.00 | 0.00 | 0.00 | 0.00 | 0.52 | 0.79 | 0.00 | 0.00 |
| 27 | 601 | 0 | 0 | 1586 | 630 | 23 | 0.00 | 0.00 | 0.00 | 0.00 | 0.00 | 0.00 | 0.50 | 0.79 | 0.00 | 0.00 |
| 28 | 181 | 0 | 0 | 556 | 195 | 9 | 0.00 | 0.00 | 0.00 | 0.00 | 0.00 | 0.00 | 0.88 | 0.95 | 0.00 | 0.00 |
| 29 | 48 | 0 | 0 | 119 | 45 | 0 | 0.00 | 0.00 | 0.00 | 0.00 | 0.00 | 0.00 | 0.61 | 0.79 | 0.00 | 0.00 |
| 30 | 10 | 0 | 0 | 23 | 8 | 0 | 0.01 | 0.02 | 0.02 | 0.02 | 0.17 | 0.17 | 0.74 | 0.86 | 0.01 | 0.02 |

**Supplementary Table 5.** Repeat unit distribution and statistical comparisons for D5S346 across six samples.

| Repeat unit length | Read count (per sample) |  |  |  |  |  | Dirichlet-multinomial test vs PBMC8 (p, padj) |  |  |  |  |  |  |  |  |  |
| --- | --- | --- | --- | --- | --- | --- | --- | --- | --- | --- | --- | --- | --- | --- | --- | --- |
|  | PBMC8 | PBMC7 | TK6 | U2OS | HCT15 | HCT116 | PBMC7_p | PBMC7_pa | TK6_p | TK6_padj | U2OS_p | U2OS_padj | HCT15_p | HCT15_pa | HCT116_p | HCT116_padj |
| 11 | 7 | 0 | 17 | 9 | 0 | 19 |  |  | 0.19 | 0.21 | 0.86 | 0.86 |  |  | 0.12 | 0.17 |
| 12 | 70 | 15 | 67 | 84 | 0 | 79 | 0.00 | 0.00 | 0.86 | 0.86 | 0.53 | 0.70 | 0.00 | 0.00 | 0.73 | 0.73 |
| 13 | 410 | 122 | 506 | 587 | 0 | 479 | 0.00 | 0.00 | 0.03 | 0.03 | 0.00 | 0.00 | 0.00 | 0.00 | 0.16 | 0.20 |
| 14 | 1760 | 807 | 2164 | 2322 | 0 | 1743 | 0.00 | 0.00 | 0.00 | 0.00 | 0.00 | 0.00 | 0.00 | 0.00 | 0.30 | 0.33 |
| 15 | 1589 | 2788 | 438 | 598 | 0 | 454 | 0.00 | 0.00 | 0.00 | 0.00 | 0.00 | 0.00 | 0.00 | 0.00 | 0.00 | 0.00 |
| 16 | 328 | 585 | 49 | 612 | 0 | 192 | 0.00 | 0.00 | 0.00 | 0.00 | 0.00 | 0.00 | 0.00 | 0.00 | 0.00 | 0.00 |
| 17 | 44 | 75 | 8 | 118 | 13 | 431 | 0.07 | 0.08 | 0.00 | 0.00 | 0.00 | 0.00 | 0.00 | 0.00 | 0.00 | 0.00 |
| 18 | 6 | 8 | 0 | 9 | 37 | 582 | 0.85 | 0.85 |  |  | 0.72 | 0.82 | 0.00 | 0.00 | 0.00 | 0.00 |
| 19 | 0 | 0 | 0 | 0 | 167 | 266 |  |  |  |  |  |  | 0.00 | 0.00 | 0.00 | 0.00 |
| 20 | 0 | 0 | 10 | 0 | 674 | 65 |  |  | 0.03 | 0.03 |  |  | 0.00 | 0.00 | 0.00 | 0.00 |
| 21 | 0 | 0 | 77 | 0 | 1486 | 19 |  |  | 0.00 | 0.00 |  |  | 0.00 | 0.00 | 0.00 | 0.00 |
| 22 | 0 | 0 | 234 | 0 | 1261 | 9 |  |  | 0.00 | 0.00 |  |  | 0.00 | 0.00 |  |  |
| 23 | 0 | 0 | 470 | 0 | 421 | 0 |  |  | 0.00 | 0.00 |  |  | 0.00 | 0.00 |  |  |
| 24 | 0 | 0 | 201 | 0 | 86 | 0 |  |  | 0.00 | 0.00 |  |  | 0.00 | 0.00 |  |  |
| 25 | 0 | 0 | 28 | 0 | 16 | 0 |  |  | 0.00 | 0.00 |  |  | 0.00 | 0.00 |  |  |

**Supplementary Table 6.**
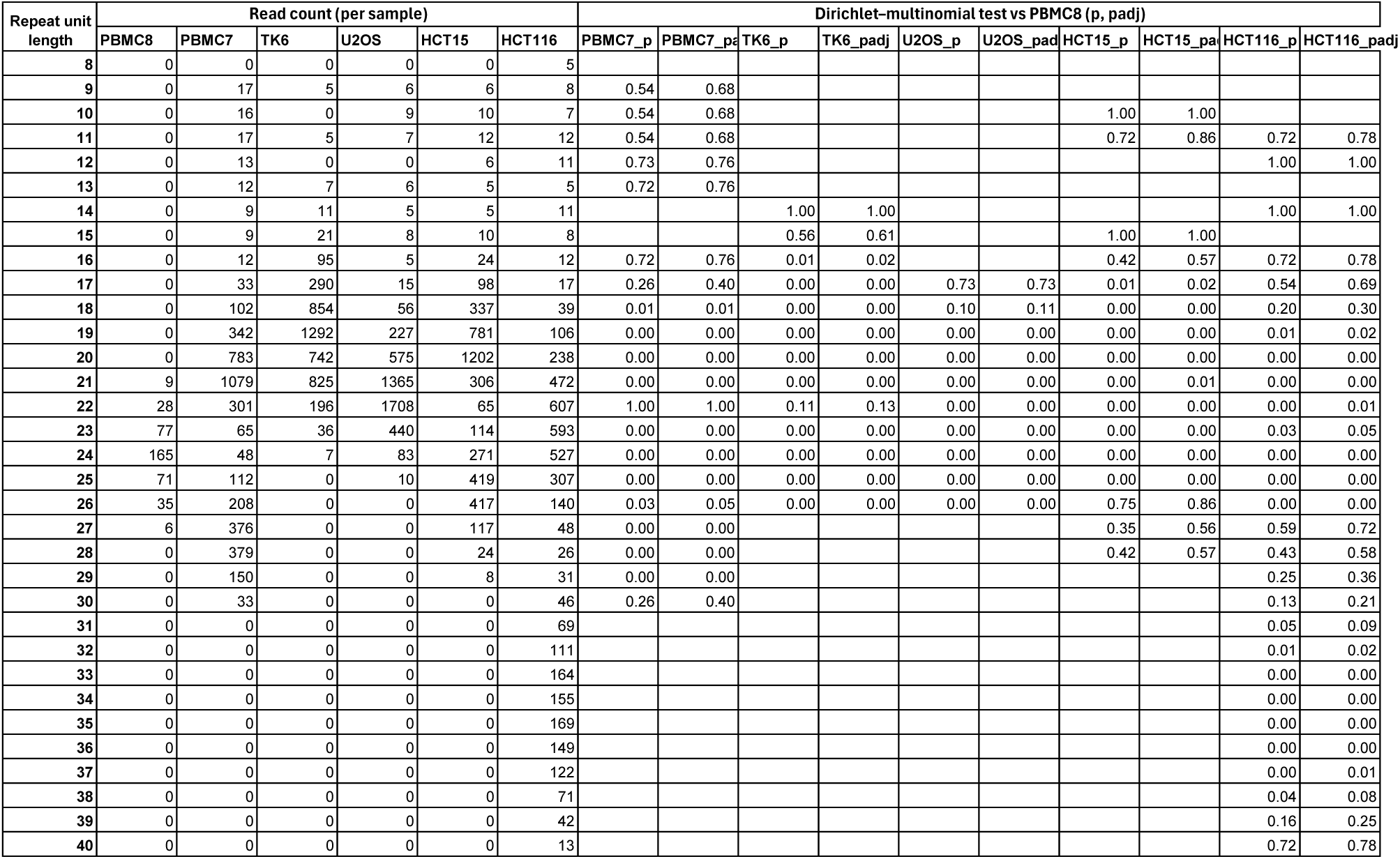
Repeat unit distribution and statistical comparisons for D17S250 across six samples.

